# A Transferable Deep Learning Approach to Fast Screen Potent Antiviral Drugs against SARS-CoV-2

**DOI:** 10.1101/2020.08.28.271569

**Authors:** Shiwei Wang, Qi Sun, Youjun Xu, Jianfeng Pei, Luhua Lai

## Abstract

The COVID-19 pandemic calls for rapid development of effective treatments. Although various drug repurpose approaches have been used to screen the FDA-approved drugs and drug candidates in clinical phases against SARS-CoV-2, the coronavirus that causes this disease, no magic bullets have been found until now. We used directed message passing neural network to first build a broad-spectrum anti-beta-coronavirus compound prediction model, which gave satisfactory predictions on newly reported active compounds against SARS-CoV-2. Then we applied transfer learning to fine-tune the model with the recently reported anti-SARS-CoV-2 compounds. The fine-tuned model was applied to screen a large compound library with 4.9 million drug-like molecules from ZINC15 database and recommended a list of potential anti-SARS-CoV-2 compounds for further experimental testing. As a proof-of-concept, we experimentally tested 7 high-scored compounds that also demonstrated good binding strength in docking study against the 3C-like protease of SARS-CoV-2 and found one novel compound that inhibited the enzyme with an IC_50_ of 37.0 μM. Our model is highly efficient and can be used to screen large compound databases with billions or more compounds to accelerate the drug discovery process for the treatment of COVID-19.

---

COVID-19 is a newly emerged infectious disease that becomes a worldwide pandemic. According to the World Health Organization (WHO) statistics, tens of millions of confirmed cases of COVID-19 and many hundreds of thousands of deaths have been reported^1^. SARS-CoV-2, a new coronavirus, has been identified to cause COVID-19^2,3^. Coronaviruses (CoVs) are a group of enveloped single stranded positive-sense RNA viruses which are able to infect many animals and humans and cause a wide range of diseases^4^. Whole genome sequencing showed that SARS-CoV-2 shares 79.6% sequence identify to SARS-CoV^5^. SARS-CoV-2 appears to have relatively high transmission rate among humans and causes severe and fatal pneumonia and other damages, threatening people at all ages, especially senior ones^6^. As the number of infections and deaths are rapidly increasing, there is an urgent call for drug and vaccine development against COVID-19.

Though immediately needed, developing new drugs within a short period of time is unpractical. Repurposing of clinically approved drugs provides a fast and effective strategy to identify antiviral drugs for immediate use. Several drugs such as remdesivir^7^, chloroquine^7,8^ and lopinavir^9^ have shown antiviral activity *in vitro* and been tested in clinical trials. FDA-approved drugs as well as those that were previously found to inhibit SARS-CoV and MERS-CoV, have been screened for their anti-SARS-CoV-2 activities^10-14^. Nine approved HIV-1 protease inhibitors were also evaluated for their anti-SARS-CoV-2 activities *in vitro* and nelfinavir was found to be active^14^. Traditional Chinese medicines provide rich resources for developing antiviral drugs. Baicalein, an ingredient isolated from *Scutellaria baicalensis* (Huangqin in Chinese), has been reported to inhibit the 3C-like protease (3CL^pro^) of SARS-CoV-2 and SARS-CoV-2 replication *in vitro*, which provides potential treatment of COVID-19^15,16^. The crystal structure of 3CL^pro^ from SARS-CoV-2 was quickly solved and docking-based virtual screening was applied. An active compound, cinanserin, which showed an IC_50_ of 125 μM for SARS-CoV-2 3CL^pro^ in the enzymatic assay, was screened out^17^.

Comparing to experimental screening and docking-based screening, deep learning based virtual screening provides a new approach in drug discovery. It generally encodes molecules into vectors and then constructs a mapping relationship from these vectors to their properties. Artificial intelligence (AI)-based virtual screening methods enable rapid search against large molecular libraries containing 10^6^∼10^9^ molecules. Stokes et al. discovered a new antibiotic with a broad-spectrum bactericidal activity by combining *in silico* predictions and experimental investigations^18^. Ton et al. applied Deep Docking model to screen all the 1.3 billion compounds from ZINC15 library and recommended the top 1,000 hits as potential SARS-CoV-2 3CL^pro^ inhibitors, though no experimental testing has been reported^19^. In the present study, we implemented a directed message passing neural network to learn the structure-activity relationship from a collection of anti-beta-CoV active and inactive compounds. Our trained model gave good predictions for the recently identified anti-SARS-CoV-2 compounds when screening the Drug Repurposing Hub library containing 6,235 FDA-approved drugs, clinical trial drugs, and pre-clinical tool compounds^20^. We then fine-tuned the model with actives and inactives against SARS-CoV-2 and applied it to screen a large compound library with 4.9 million drug-like molecules from ZINC15 database^21^. We suggested a list of potential anti-SARS-CoV-2 compounds for further experimental testing. As a proof-of-concept, we experimentally tested the activities of 7 molecules with high prediction scores and good binding affinities from docking studies against 3CL^pro^ of SARS-CoV-2 and found one active compound with an IC_50_ of 37.0 μM.

## Material and Methods

### Data

Training dataset is essential for deep learning methods. In order to train a robust model that can predict new antiviral drugs against SARS-CoV-2, an ideal training set should contain sufficient positive and negative compounds for SARS-CoV-2. Unfortunately, SARS-CoV-2 is a newly emerged coronavirus, and only limited information is available now. SARS-CoV-2, as well as HCoV-OC43, SARS-CoV and MERS-CoV, belongs to beta-coronaviruses^3,22^. They share a high degree of conservation in essential functional proteins, including the 3CL^pro^, the RNA-dependent RNA polymerase, the RNA helicase, etc.^23^. For example, the 3CL^pro^ in SARS-CoV and SARS-CoV-2 share a sequence identity of 96.1%, indicating that these CoVs share potential targets for broad-spectrum anti-CoV drugs. Potent MERS-CoV inhibitors identified by screening an FDA-approved drug library also inhibit the replication of SARS-CoV and HCoV-229E^24^. Shen et al. found seven broad-spectrum antiviral inhibitors through a high-throughput screening of a 2,000-compound library against HCoV-OC43^23^. These studies provide a list of antivirals for beta-CoVs that can be used to train a model for screening SARS-CoV-2 antiviral candidates.

We collected a set of inhibitors against HCoV-OC43, SARS-CoV and MERS-CoV from literatures with a cutoff of EC_50_ < 10 μM and selective index > 10^22,23,25-27^. All the inhibitors were identified by screening libraries including FDA-approved drugs and pharmacologically active compounds. After applying the cutoff filter, 90 compounds were selected as antivirals and each of them can inhibit at least one of the three CoVs. The remaining compounds were regarded as negative data. This primary training dataset (Training Set 1) containing 90 positives and 1,862 negatives was used to train the deep learning classification model for screening anti-beta-coronavirus compounds.

We also constructed an independent data set containing a collection of experimentally tested active and inactive molecules against SARS-CoV-2^10-14^. We labelled these compounds by their activity against SARS-CoV-2 according to a cutoff of EC_50_ < 50 μM, resulting in 70 actives and 84 inactives (Fine-tuning Set 1). We applied this SARS-CoV-2-specific dataset to train the SARS-CoV-2-specific antiviral prediction model.

The Drug Repurposing Hub is a curated and annotated collection of FDA-approved drugs, clinical trial drug candidates, and pre-clinical compounds with a companion information resource. We applied our model to this library to identify potential antiviral molecules. Compounds overlapping with the training dataset were removed and the rest compounds were used to screen potential antivirals.

ZINC15 is a free database designed for virtual screening, containing ∼1.5 billion molecules^21^. We extracted a subset database containing ∼4.9 million molecules that are drug-like and in stock. Virtual screening was applied to this library to discover potential antiviral molecules.

### Model

In this work, we developed a series of COVID19-related Virtual Screening (COVIDVS) models, which implemented a directed-message passing deep neural network model based on Chemprop that has been used to predict molecular properties directly from the graph structure of molecules^28^. Chemprop model takes molecular SMILES as input and converts it to a graph representation internally. Atoms and bonds are regarded as graph nodes and edges, respectively. A related feature vector is assigned to each atom and bond, then the model is trained to encode information about neighboring atoms and bonds via a directed bond-based message passing approach. Finally, a vector representing the whole molecule is generated by combining all the bond messages. An additional vector containing molecular features computed by RDKit^29^ is concatenated to molecular representation to avoid overfitting. The Ensemble method has been shown to be able to improve the performance of machine learning models, which is accomplished by training the same model architecture several times with different random initial weights and then averaging the results^30^. Here we applied ensemble method to construct our COVIDVS model for better performance.

Transfer learning (TL) is an AI technology that can be applied to resolve problems of data scarcity by leveraging existing knowledge from other related tasks to a specific task with low data^31^. Transfer learning have achieved success on low data tasks in many fields including computer vision^32^, natural language processing^33,34^ and drug discovery^35,36^. In the present study, we implemented fine-tuning technique, which is one of the most commonly used transfer learning techniques, to deal with the data scarcity problem for anti-SARS-CoV-2 prediction model.

## Results

### The broad-spectrum anti-beta-coronavirus compound prediction model

We used the Training Set 1, which consists of 1,952 compounds labeled by their activities against SARS-CoV, MERS-CoV or HCoV-OC43 to train a general classification model for anti-beta-CoV activity. We carried out Bayesian optimization to define the hyperparameters. In order to evaluate the performance of model, we trained models on the training data from each of the 10 different random splits of Training Set 1, each with 80% training data, 10% validation data and 10% test data, resulting in an average of receiver operating characteristic curve-area under the curve (ROC-AUC) of 0.96 on the training data and 0.83 on the testing data (Extended Data Fig. 1). We then constructed an ensemble of 5, 10, 20 models respectively and tested their performance on an independent testing set, which was constructed by removing the overlapping compounds within Fine-tuning Set 1 and Training Set 1 from Fine-tuning Set 1. The independent testing set (named Test Set 1) consists of 33 active molecules and 38 inactive molecules. The ensemble of 20 models achieved the best performance with a ROC-AUC of 0.89 on Test Set 1 (see Fig. 1a and Extended Data Fig. 2), indicating that this model can efficaciously discriminate actives and inactives for SARS-CoV-2. Therefore, we selected the ensemble of 20 models (COVIDVS-1) for further prediction.

**Fig. 1.**
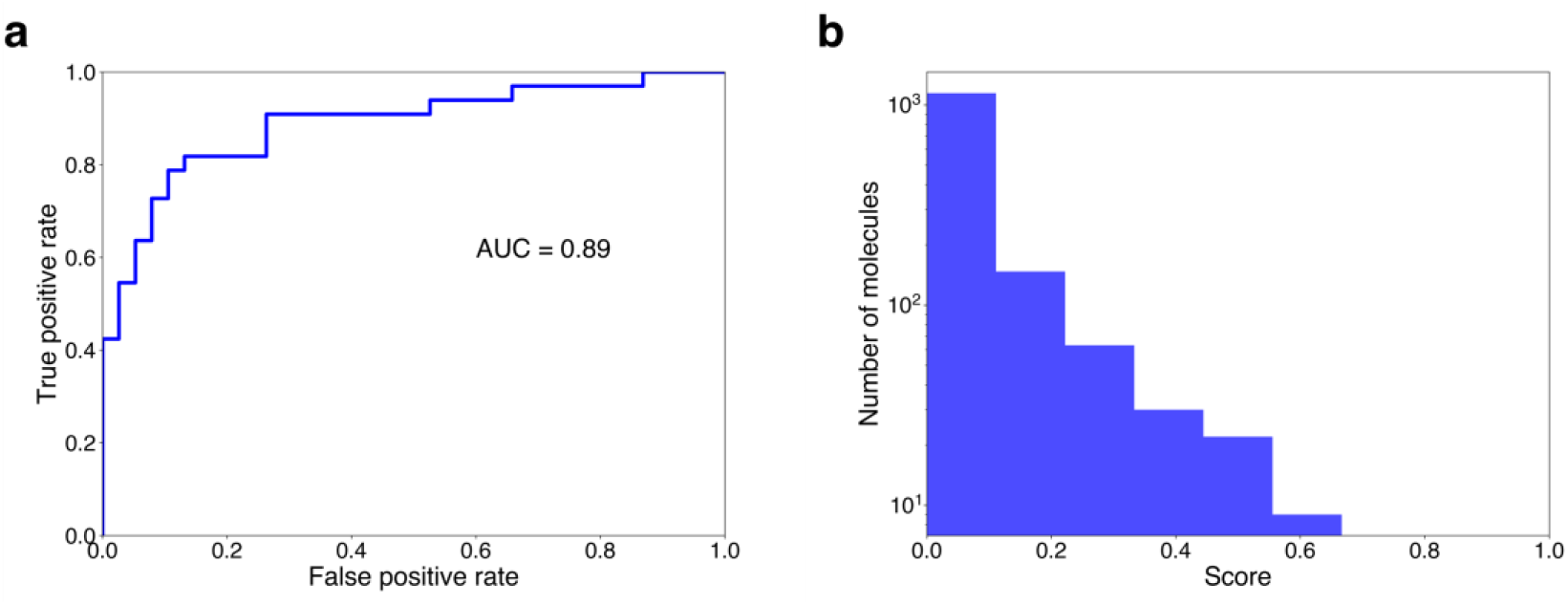
The performance and prediction results of COVIDVS-1 model. **a**, ROC curve showing the performance of COVIDVS-1 on Test Set 1. **b**, Histogram showing the distribution of predicted scores of the launched drugs library extracted from Drug Repurposing Hub. Molecules that are in Training Set 1 have been removed.

We applied COVIDVS-1 to predict the anti-SARS-CoV-2 activity of compounds in a library containing 1,417 launched drugs extracted from the Drug Repurposing Hub. Fig. 1b gives the distribution of the predicted scores. Most of the launch drugs (89.3%) have scores less than 0.2. Among the 70 top-ranking (5%) drugs, 6 have been reported to be active against SARS-CoV-2 (Table 1). Ceritinib (also named LDK378), a drug that is used for the treatment of non-small-cell lung cancer, ranks at position 11 and was reported to inhibit the replication of SARS-CoV-2 with an IC_50_ of 2.86 μM^10^. Terconazole, an antifungal drug that ranks at position 24, showed an IC_50_ of 11.92 μM ^11^. Osimertinib, an anti-cancer drug that was used to treat non-small-cell lung carcinomas with a specific mutation, ranks at position 35 and has been shown to be active against SARS-CoV-2 with an IC_50_ of 3.26 μM^10^. Ritonavir, an antiretroviral medication used along with other medications to treat AIDS, showed 8.63 μM of EC_50_ and ranks at position 42^14^. Abemaciclib, a drug for the treatment of breast cancers that showed potency against SARS-CoV-2 with an IC_50_ of 6.62 μM, ranks at position 46^10^. Indinavir, a protease inhibitor used as a component of highly active antiretroviral therapy to treat AIDS, ranks at position 60 with a reported anti-SARS-CoV-2 EC_50_ of 59 μM^14^. These results demonstrated that COVIDVS-1 can successfully screen out potential antiviral drugs against SARS-CoV-2, even though the active compounds in the training set were only tested on beta coronaviruses other than SARS-CoV-2. We analyzed the chemical structure similarity between the 6 drugs and the active compounds in the Training Set 1 by Tanimoto similarity coefficient with Morgan Fingerprint (Calculated by RDKit). All these 6 drug molecules showed different chemical structures with maximum similarity < 0.4, indicating that the model can identify potential candidates with novel structures.

**Table 1.** COVIDVS-1 predicted ranking positions of six launched drugs reported to have antiviral activity against SARS-CoV-2 ^10,11,14^ among the 1417 launched drugs extracted from the Drug Repurposing Hub.

| Name | Experimental activity (EC<sub>50</sub>) | Ranking position |
| --- | --- | --- |
| Ceritinib | 2.86 µM | 11 |
| Terconazole | 11.92 µM | 24 |
| Osimertinib | 3.26 µM | 35 |
| Ritonavir | 8.63 µM | 42 |
| Abemaciclib | 6.62 µM | 46 |
| Indinavir | 59 µM | 60 |

### Development of anti-SARS-CoV-2 compound prediction model

Fine-tuning Set 1 contains only 154 data, which is obviously not sufficient to allow us to train a new model from scratch. Instead, we used the Fine-tuning Set 1 to fine-tune our COVIDVS-1 model, resulting in the second generation model, COVIDVS-2. COVIDVS-2 contains information from both Training Set 1 and Fine-tuning Set 1. We added all data in the Fine-tuning Set 1 to the Training Set 1 to build the Training Set 2. Molecules already existed in the Training Set 1 were relabeled according to their activity against SARS-CoV-2 and molecules not presented in the Training Set 1 were directly added. The Training Set 2 contains 133 positive data and 1,890 negative data. We analyzed the chemical space distribution of the Training Set 2 and data from the Drug Repurposing Hub using the t-Distributed stochastic neighbor embedding (t-SNE) dimension reduction method. Tanimoto similarity was utilized to quantify the chemical distance. The positive data in the training set largely overlaps with the data from Drug Repurposing Hub in chemical space (Fig. 2a). We then used this model to screen the full Drug Repurposing Hub dataset. The distribution of the predicted scores is given in Fig. 2b. There are 280 molecules with score > 0.8 and 55 molecules with score > 0.9. About half of the top 55 molecules were reported kinase inhibitors, demonstrating the potential of using kinase inhibitors as anti-SARS-CoV-2 drugs. Six anaplastic lymphoma kinase (ALK) tyrosine kinase receptor inhibitors, three cyclin-dependent kinase (CDK) inhibitors and eight epidermal growth factor receptor (EGFR) tyrosine kinase inhibitors were enriched in the top 55 list, which have the same targets to Ceritinib, Abemaciclib and Osimertinib that are active on SARS-CoV-2, respectively. Compared to the ∼40 known targets of the 55 molecules and the ∼60 known targets of active molecules in Training Set 2, only 8 targets are the same, demonstrating that the molecular targets of predicted results were not constrained by the training set. We listed all the 55 molecules with score > 0.9 in Supplementary Table S1 and grouped them according to their clinical study states. While preparing this manuscript, we noticed a newly reported work that carried out a mass spectrometry-based phosphoproteomics survey of SARS-CoV-2 early infection. Dramatic rewiring of phosphorylation on host and viral proteins and alter activities of kinases were observed during the SARS-CoV-2 infection, making kinases to be ideal drug targets^37^.

**Fig. 2.**
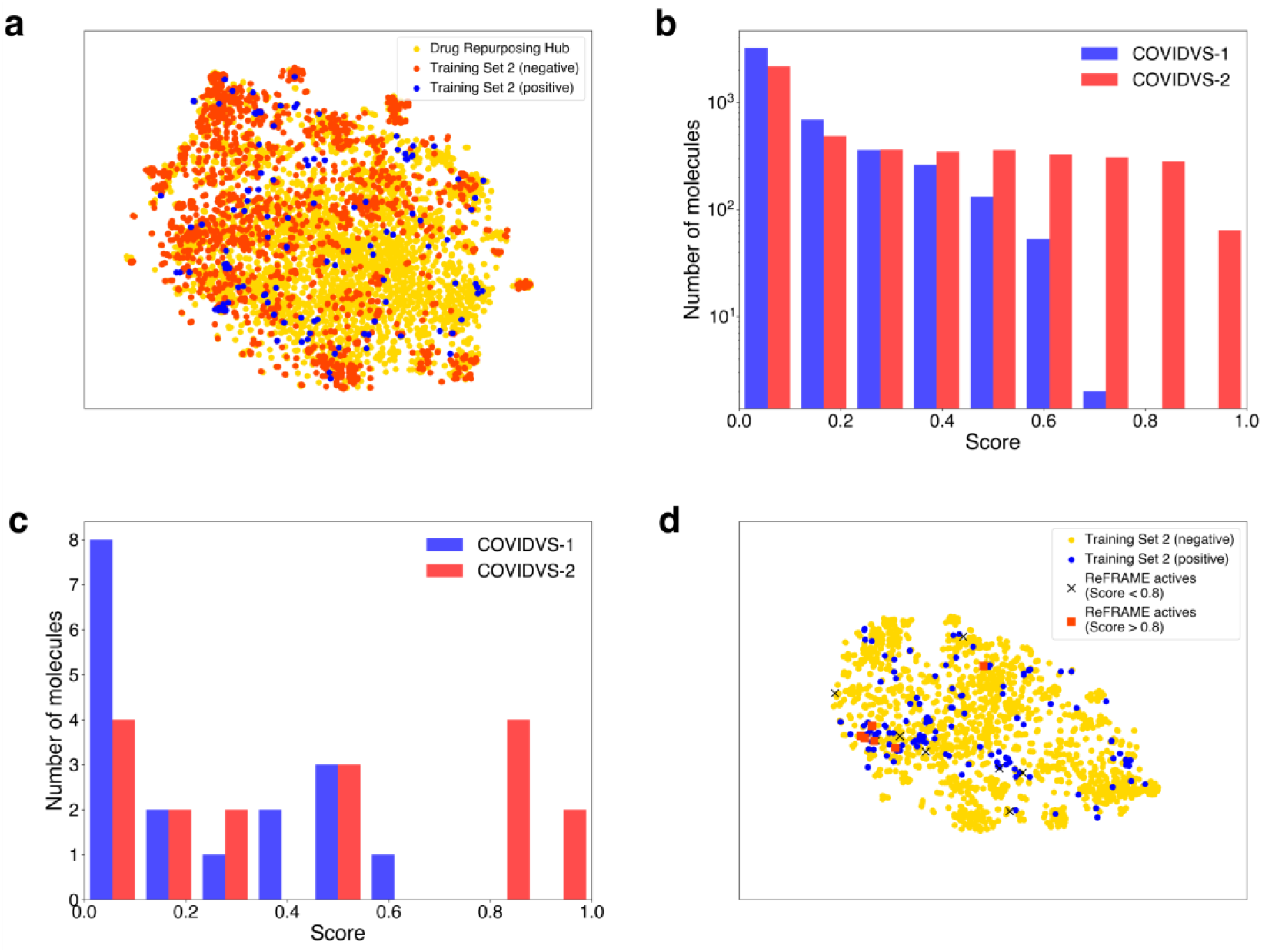
Comparison the prediction results between COVIDVS-1 and COVIDVS-2 (b, c) and the data distribution Training Set 2, Drug Repurposing Hub and ReFRAME actives (a, d). **a**, t-SNE of all molecules from the Training Set 2 (red: negative data, blue: positive data) and Drug Repurposing Hub (gold). **b**, Histogram showing the difference between prediction results of COVIDVS-1 and COVIDVS-2 on Drug Repurposing Hub. **c**, Distribution of scores for 17 ReFRAME actives predicted with COVIDVS-1 and COVIDVS-2, respectively. **d**, t-SNE of all molecules from the Training Set 2 (blue: positive data, gold: negative data) and ReFRAME actives (black cross: 11 molecules with score < 0.8, red squares: 6 molecules with score > 0.8).

A recent study experimentally screened the ReFRAME library, which collects a large number of clinical-phase or FDA-approved drugs, against SARS-CoV-2 and found 20 active compounds^38,39^. Among the 20 active compounds, 3 were already included in our Training Set 1 and Fine-tuning Set 1. We used the 17 newly identified active compounds (hereafter referred as ReFRAME actives) to evaluate our COVIDVS-1 and COVIDVS-2. We predicted the ReFRAME actives with COVIDVS-2 and 6 of them have scores > 0.8. Compound KW 8232 (EC_50_∼1.2 μM) has a score of 0.94, which is above most of the 4,711 Drug Repurposing Hub molecules. This suggests that our model can successfully discover novel antiviral drugs against SARS-CoV-2. We also compared the performance of COVIDVS-2 and COVIDVS-1 on ReFRAME actives (Fig. 2c). Among the 17 compounds, six got predicted scores > 0.8 by COVIDVS-2, while no molecule got predicted score > 0.8 by COVIDVS-1. Of course, higher scores may not guarantee true activity. We mixed these 17 molecules into the 4,711 molecules from the drug Repurposing Hub and ranked all of them by their predicted scores. The top-2 compounds among the 17 ranked in 28th and 58^th^ among all the 4,728 compounds when predicted with COVIDVS-1, while the ranking raised to 4th and 29th when predicted with COVIDVS-2. We posted these ReFRAME actives onto the t-SNE plot of the Training Set 2. Although all the 6 compounds with good predictions are close to active compounds in the training set, some of the 11 compounds with predicted scores less than 0.8 are relatively far from the active compounds in the Training Set 2 (Fig. 2d). This demonstrates that the diversity of active compounds limited the model performance, which can be improved by increasing the number and chemical diversity of active compounds in the training set data.

### Screening ZINC database to identify novel anti-SARS-CoV-2 compounds

Though several FDA-approved drugs have shown anti-SARS-CoV-2 activities using drug repurposing approaches, none of them were highly effective in clinical trials. Highly effective novel anti-SARS-CoV-2 drugs need to be developed. Deep learning models can be easily applied to deal with big data, which allows us to screen large chemical libraries. We subsequently applied our method to screen ZINC15 database. We added all the ReFRAME actives to the Fine-tuning Set 1 to construct the Fine-tuning Set 2. We then fine-tuned the COVIDVS-1 model with the Fine-tuning Set 2 to derive the third-generation model, COVIDVS-3.

We applied COVIDVS-3 to screen the 4.9 million drug-like molecules selected from ZINC15. This screen run was finished within 6 hours using 200 CPUs for feature generation and 4 NVIDIA GPUs for prediction, which can be easily speeded up. We ranked all the 4.9 million molecules by their predicted scores. This gave 3,641 molecules with score > 0.9, and 94.6% of them have the maximum similarity < 0.4 to positive data of Training Set 3. In order to understand the structure distribution relationship of the high-score molecules, we analyzed the chemical space distribution of the 3641 molecules and the positive data in Training Set 3. A Density-Based Spatial Clustering of Applications with Noise (DBSCAN) method^40,41^ was performed to cluster the 3,641 ZINC molecules with score > 0.9 (See Methods section for details). There are 46 clusters that have at least 10 molecules, and 8 clusters with at least 50 molecules. Fig. 3 shows the clustering results on t-SNE plot, as well as the distribution of positive data of Training Set 3, revealing that the predicted compounds locate in different chemical space, showing the diversity of results. For each of the 8 clusters with at least 50 molecules, we selected one molecule with the best score as representative compounds (shown in **Fig. 4**). The top 100 molecules with best prediction scores and representative molecules from the 46 clusters were given in Supplementary Table S2. We suggest that these compounds be tested for their anti-SARS-CoV-2 activities in future experimental studies. Our method can be easily applied to screen other large library with millions, even billions of compounds.

**Fig. 3.**
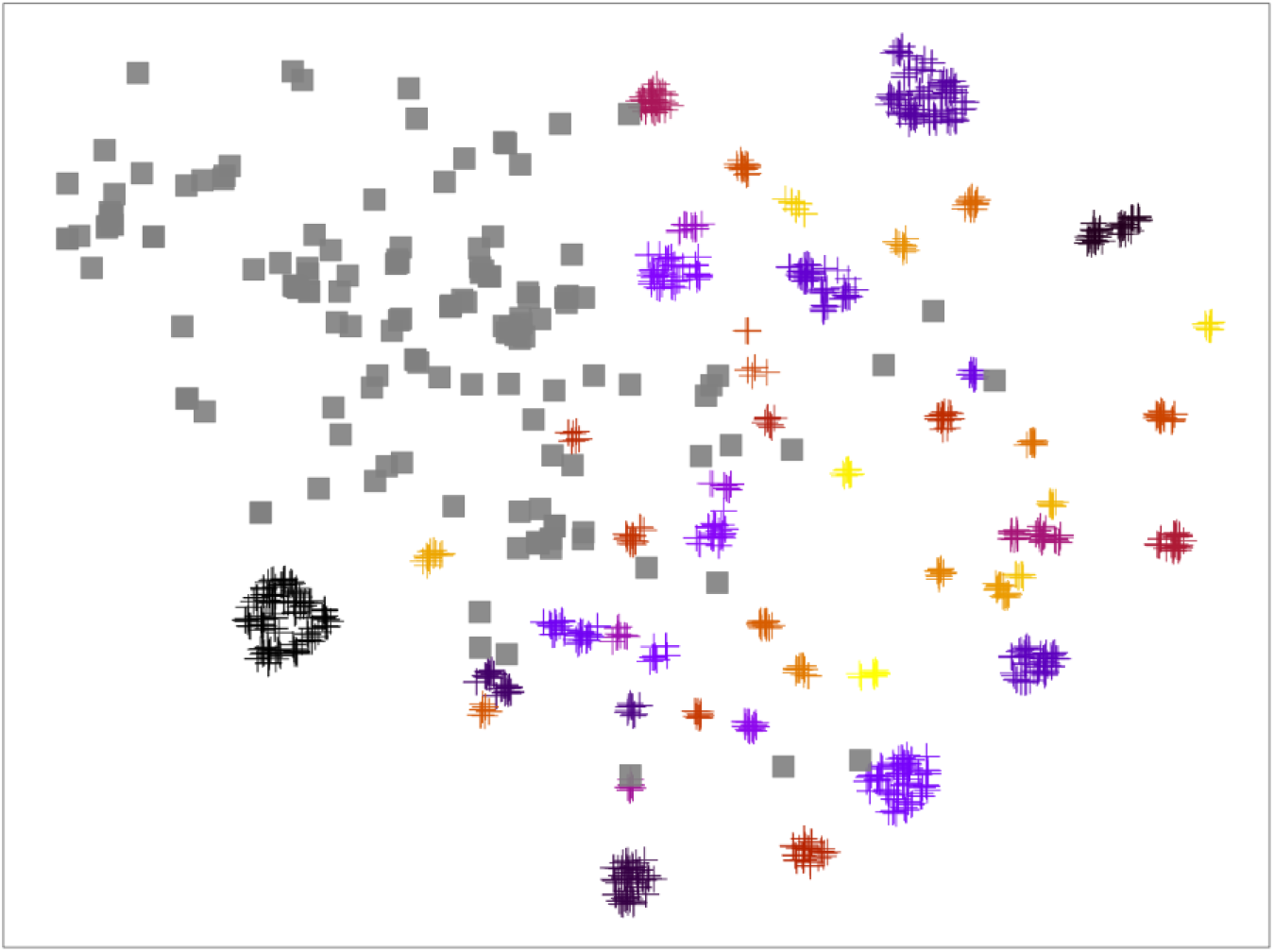
Clustering results of high-score ZINC15 molecules. Molecules from positive data of the Training Set 3 and the predicted ZINC15 molecules with score > 0.9 were plotted on t-SNE distribution plot. Grey squares represent the positive data of the Training Set 3. Plus markers with different colors represent different clusters. Noisy samples which are far away from any clusters are not plotted for clarity.

**Fig. 4.**
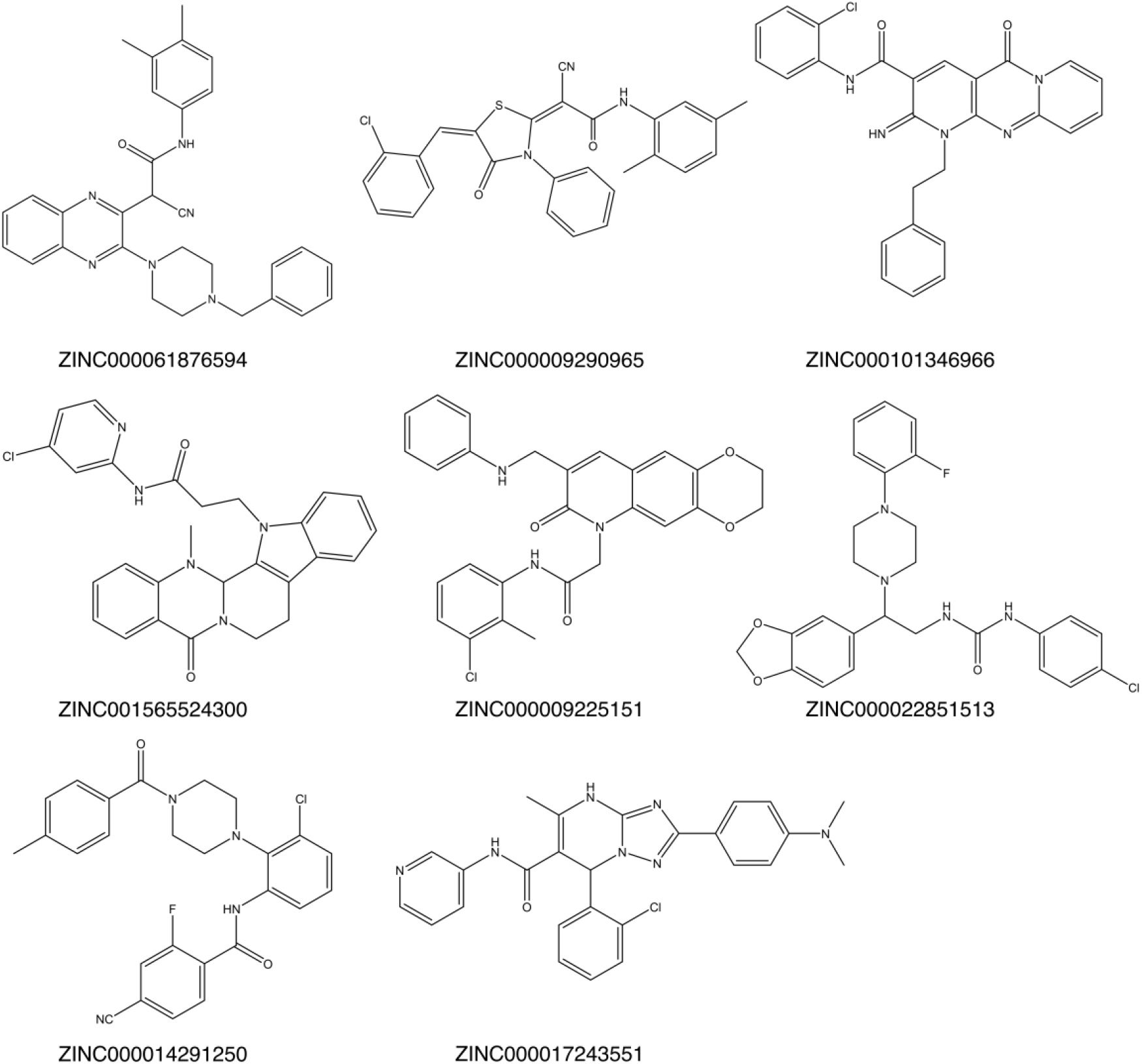
2D structures of the eight ZINC15 molecules with best score from each of clusters that have at least 50 molecules.

### Identifying novel SARS-CoV-2 3C-like protease inhibitors

We have applied our COVIDVS model to screen ZINC15 database and predicted a set of potential antiviral molecules, which may act on different targets, including the SARS-CoV-2 3CL^pro^. 3CL^pro^ plays an important role in mediating viral replication and transcription^42^. The sequence identity of 96.1% between 3CL^pro^ in SARS-CoV and SARS-CoV-2 makes it an ideal target for developing broad-spectrum anti-CoV drugs. In order to further screen 3CL^pro^ inhibitors from the prediction results, we performed molecular docking using Autodock Vina software^43^. The structure of SARS-CoV-2 3CL^pro^ (PDB ID 6LU7)^17^ and candidate molecules were prepared with AutodockTools^44^. All the 3,641 ZINC15 molecules with prediction score > 0.9 from the previous section were subjected to docking. Their docking scores ranges from −10.5 to −6.3 kcal/mol. From the top 40 results, we selected 7 easily purchasable compounds to experimentally evaluate their activities.

We purchased these 7 compounds and tested their SARS-CoV-2 3CL^pro^ inhibition activity (see Methods section for experimental details). Among all the 7 compounds tested, ZINC000017053528 showed strong inhibition at 50 μM (Supplementary Table S3) with an IC_50_ of 37.0 μM (**Fig. 5**). To the best of our knowledge, no bioactivity of this molecule has been reported before. We calculated the 2D structure similarity between the active compound and the 405 reported SARS-CoV 3CL^pro^ inhibitors from PubChem AID1706 assay^45^ with ECFP4 fingerprint^46^. All the known active compounds have the similarity less than 0.4, demonstrating that this newly discovered active compound has a novel chemical structure compared to known inhibitors.

**Fig. 5.**
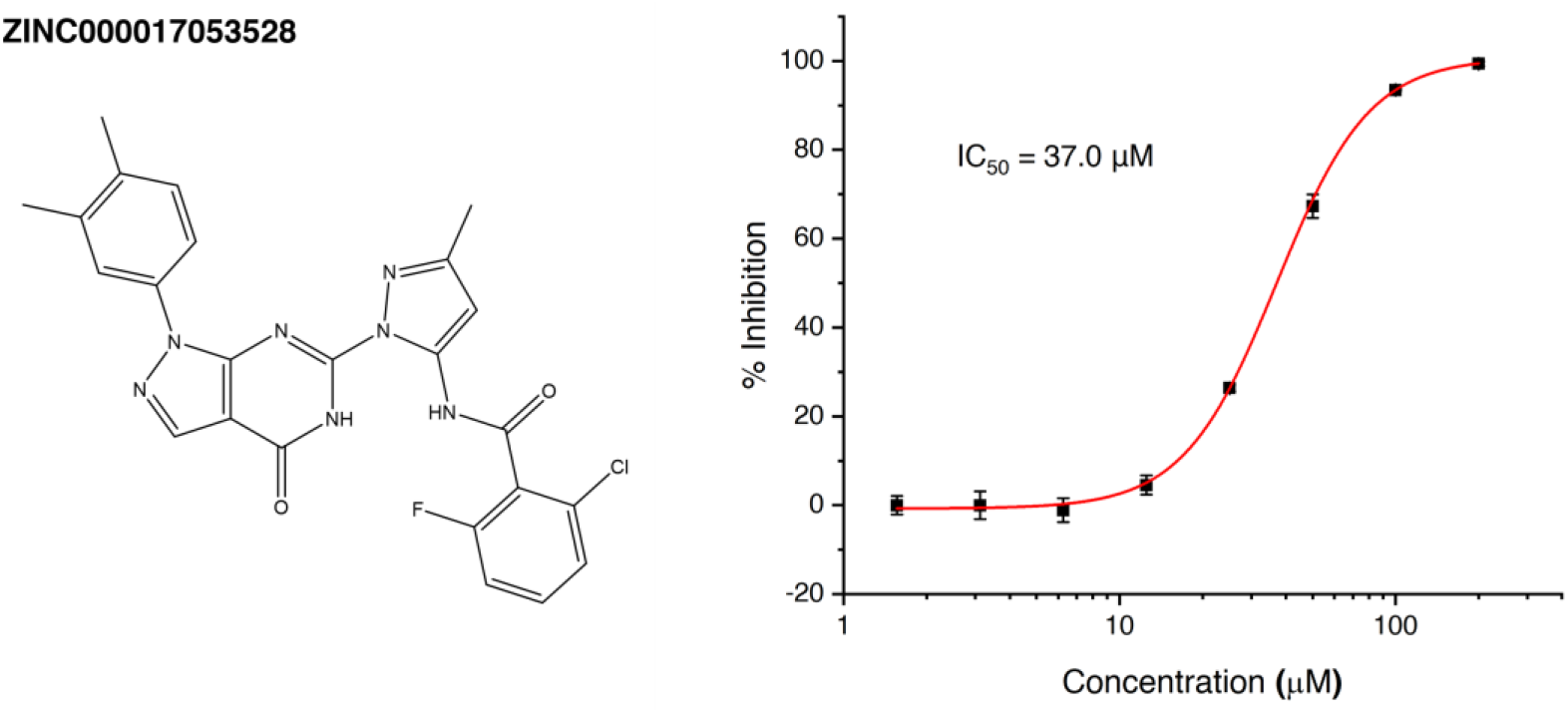
The chemical structure (left) and the *in vitro* anti-SARS-CoV-2 3CL^pro^ activity (right) of molecule ZINC000017053528.

## Discussion

We built a directed-message passing neural network model to fast screen potential drugs against SARS-CoV-2. Our model was firstly trained with a collection of broad-spectrum anti-beta-coronavirus compounds and then migrated the extracted knowledge to anti-SARS-CoV-2 prediction model through transfer learning. The ensemble technique helped to improve the performance of our COVIDVS model, however, it also increased the computational cost proportionally. Therefore, the balance between performance and cost should be taken into consideration. Although the ensemble of 20 models showed the best performance, we have demonstrated that the ensemble of 5 or 10 models are quite effective to improve the model’s performance (Extended Data Fig. 2) and can be applied when screening ultra-large compound libraries to reduce the computational cost.

As a data-driven method, the performance of deep learning model heavily depends on the training data, which should include enough known molecules with and without antiviral activities in this case. In order to overcome the lack of data for SARS-CoV-2, we collected data from three other coronaviruses, SARS-CoV, MERS-CoV and HCoV-OC43, which all belong to beta-coronaviruse group like SARS-CoV-2, and have been widely studied. Molecules screened for one of the CoVs often showed broad-spectrum anti-CoVs activities, indicating that these data can help us to discovery new antiviral compounds for SARS-CoV-2. We have demonstrated that the broad-spectrum antiviral prediction model COVIDVS-1 trained with Training Set 1 can successfully screen potential active molecules against SARS-CoV-2 in the top list of prediction results. The newly discovered active compounds targeting SARS-CoV-2, as well as the other three coronaviruses, in turn, can be used to enhance the performance of the broad-spectrum model, which can be applied to screen potential broad-spectrum drugs for newly emerged coronaviruses in the future. Fine-tuning technique provides another way to improve the model’s performance, resulting in a task-specific model. It is expected that more data will further improve the prediction ability of transfer learning-based model, which can be achieved by iterating the fine-tuning, model prediction and experimental estimation process. This strategy can be easily achieved when we are facing an interesting target lacking enough data.

Due to experimental limitations, we were unable to test our predicted compounds on SARS-CoV-2 directly in our own laboratory. As an alternative experimental validation, we used our COVIDVS prediction together with protein-ligand docking to screen for potential SARS-CoV-2 3CL^pro^ inhibitors. We performed docking of the 3,641 top-ranking compounds instead of the 4.9 million drug-like molecules from ZINC15 database and identified a new SARS-CoV-2 3CL^pro^ inhibitor with novel chemical scaffold, which can be further optimized. Although many 3CL^pro^ inhibitors have been reported, most of them only showed activity in *in vitro* enzyme assay. As our COVIDVS models were trained with antiviral activity data, compounds with *in vitro* 3CL^pro^ inhibition activity and good COVIDVS prediction scores may have high probability of anti-viral activity. Similar to COVIDVS-2 and 3, a target-specific models for 3CL^pro^ can be trained by fine-tuning COVIDVS-1 with known 3CL^pro^ inhibitors and non-inhibitors, which is expected to increase the success rate of prediction.

## Conclusion

COVID-19 remains as a global pandemic that is waiting for effective vaccines and drugs. A number of FDA-approved drugs and clinical-phase molecules are being tested in clinical trials. However, no magic bullets have been found yet. More efforts are necessary to identify safe and efficacious therapeutic solutions for COVID-19 and emerging CoV related diseases in the future. Here we present a method that can fast discover potent antiviral drugs by screening drug repurposing library and other large virtual screening libraries *in silico*, which can largely reduce the number of compounds that need to be experimentally tested. All of the Training Set, Fine-tuning Set and Test Set were provided in Supplementary Table S4. The datasets and models can also be found at https://github.com/pkuwangsw/COVIDVS.

## Methods

### Hyperparameters

The Chemprop model includes 2 modules. The first module is a molecular encoder based on the message passing neural network (MPNN) and the second module is a feed-forward neural network (FFN). In our work, we set the depth of MPNN to 6 and the depth of FFN to 3. The hidden units of FFN layers are set to 1500 and all the dropout values were set to 0.3. The hyperparameters were defined with Bayesian optimization.

### Model training and fine-tuning

The initial model COVIDVS-1 was trained with default parameters except the hyperparameters mentioned above. COVIDVS-2 and COVIDVS-3 were trained by fine-tuning COVIDVS-1. Because only 154 data are available for fine-tuning, we frozen the weights of MPNN layers in the fine-tuning process to reduce the parameter space. Besides, we fine-tuned each of the 20 models in COVIDVS-1 and merged them as COVIDVS-2 and COVIDVS-3 models.

### Ensemble method

We constructed ensemble models by training multiple models separately and averaging their prediction results. We compared the performance of COVIDVS models with an ensemble of 1, 5, 10 and 20 models. Each ensemble model was trained with Training Set 1 and tested on Test Set 1. The ROC-AUC values were calculated to evaluate the model performance. We tested each type of models three times and the results were listed in Extended Data Fig. 2. Comparing with using one single model, an ensemble of multiple models can effectively increase the model’s performance and confidence. According to the results, we selected to ensemble 20 models in our work.

### Clustering

We applied Density-Based Spatial Clustering of Applications with Noise (DBSCAN) method^40,41^ to cluster the 3,641 ZINC molecules with score > 0.9. This method can find core samples of high density and expands clusters from them. The maximum distance between two samples for one to be considered as in the neighborhood of the other is set to 0.3 and the minimal number of samples in a neighborhood for a point to be considered as a core point is set to 10. The distance of two molecules was defined with Tanimoto Similarity. Clustering process was performed with Python 3.7 and scikit-learn’s default parameters except those mentioned before.

### Protein expression and purification

The gene encoding SARS-CoV-2 3CL^pro^ with *Escherichia coli* codon usage was synthesized by Hienzyme Biotech. The recombinant SARS-CoV-2 3CL^pro^ was expressed and purified as previously described^15^.

### The inhibition assay of SARS-CoV-2 3CL^pro^ activity

A fluorescent substrate Dabcyl-KTSAVLQSGFRKM-Edans (GL Biochemistry Ltd) and buffer composed of 40 mM PBS, 100 mM NaCl, 1 mM EDTA, 0.1% Triton X-100, pH 7.3 was used for the inhibition assay. Stock solutions of the inhibitor were prepared with 100% DMSO. The measurement was carried out with 0.5 µM protease, 20 µM substrate and DMSO or inhibitor at various concentrations. The fluorescence signal generated by the cleavage of the substrate was monitored for 20 min at an emission wavelength of 460 nm with excitation at 360 nm, using a plate reader (Synergy, Biotek). The percent of inhibition was calculated by V_i_/V_0_, where V_0_ and V_i_ represent the mean reaction rate of the enzyme incubated with DMSO or compounds. IC_50_ was fitted with Hill1 function.

## Acknowledgments

This work was supported in part by the Ministry of Science and Technology of China (2016YFA0502303), the National Natural Science Foundation of China (21633001) and the Fundamental Research Funds for the Central Universities of China. We would like to thank Dr. Xiaobing Deng, Dr. Weilin Zhang, Dr. Hao Liang, Weixin Xie and Hongyi Wang for collecting and sharing anti-beta-CoV actives/inactives. We would like to thank Chenjing Cai for her help in transfer learning technique. Part of the computation and analysis was performed on the High Performance Computing Platform of the Peking-Tsinghua Center for Life Sciences, Peking University.

## Author contributions

J.P and L.L conceived and supervised the research. S.W and Y.X constructed the deep learning model and perform computational prediction and analysis. Q.S designed and performed the validation experiments. All authors contributed to the interpretation of results. All authors wrote and approved the manuscript.

---

**Extended Data Fig. 1.**
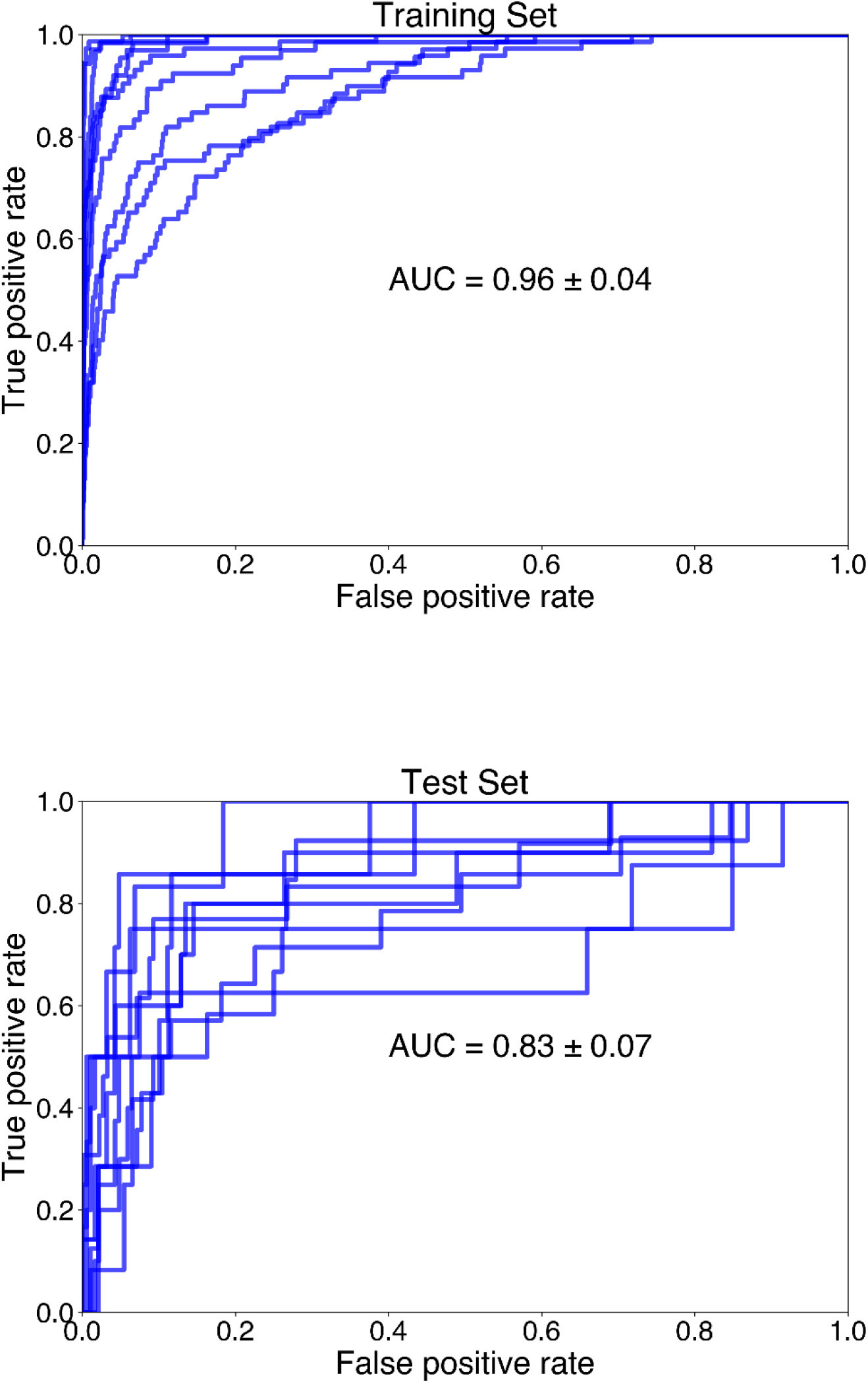
ROC curves evaluating the model’s performance on the training set (top) and test set (bottom). We trained 10 models with different random split of Training Set 1 to evaluate the model’s stability. The AUC values are the average of ten models.

**Extended Data Fig. 2.**
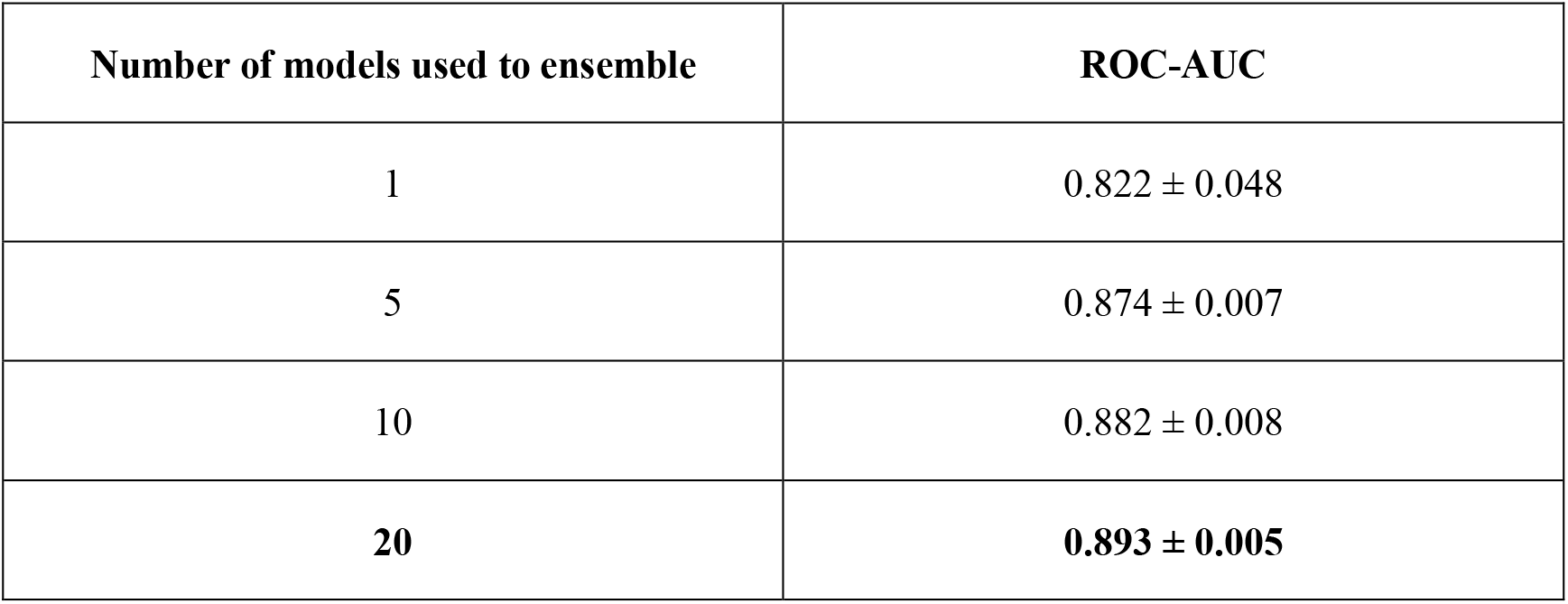
Comparison of model performance by ensembling different number of models.

